# Sensitivity of the timing of *Drosophila* pupal wing morphogenesis to external perturbations

**DOI:** 10.1101/2023.09.22.558969

**Authors:** Romina Piscitello-Gómez, Ali Mahmoud, Natalie A Dye, Suzanne Eaton

## Abstract

The morphogenesis of the pupal wing of *Drosophila melanogaster* is a powerful model for understanding the physical mechanisms of tissue morphogenesis. During this process, the wing undergoes a dramatic reshaping where the hinge region contracts and pulls on the blade region, triggering a dynamic pattern of cell elongation changes, cell divisions, and cell neighborhood changes Etournay et al. (2015). Over our several year study of this process, we noticed that sometime after the year 2016, the onset of morphogenesis became consistently delayed by ∼6 h compared to previous years. Thus, the peak of cell elongation was now observed at 28 hours after puparium formation (hAPF) rather than at 20-22 hAPF. We show here that this delay is independent of genetic background and that the speed of the overall process is relatively unchanged, implying that an environmental factor causes a shift in the onset of the process relative to the start of pupariation. We then investigated how various environmental factors influence the timing of pupal wing morphogenesis, using the dynamics of cell elongation as a proxy. We find that imaging conditions, temperature, diet, *Wolbachia* infection, and the light/dark cycle have relatively little effect. Preliminary data show that the spectra of light used to rear the flies affects cell elongation dynamics, although the delay was not completely reverted and more experiments are needed to know precisely how this variable affects morphogenesis. In sum, our work suggests the presence of a gate controlling the onset of pupal wing morphogenesis that is influenced by a specific, yet unclear, environmental factor. Once the gate is released, pupal wing morphogenesis occurs at a fairly constant rate, remarkably robust to many environmental variables.

## 1 Introduction

Tissue morphogenesis requires a tight control of cellular behaviors in space and time. Here we study the reshaping of the pupal wing epithelium of *Drosophila melanogaster*. The pupal wing undergoes a dramatic shape change over the course of approximately 16 hours. The proximal wing hinge region contracts and reshapes the wing blade, which narrows along the anterior-posterior (AP) axis and elongates along the proximal-distal (PD) axis (Etournay et al., 2015). The change in wing shape is the result of cell behaviors including cell divisions, cell neighbor changes, and cell shape changes.

Our group has focused on understanding the interplay between tissue mechanics and cell polarity during pupal wing morphogenesis. We showed that both cell rearrangements and cell shape changes contribute to tissue deformation in the pupal wing (Etournay et al., 2015). At the beginning of pupal wing morphogenesis, cell shapes are isotropic and they become anisotropic as hinge contraction proceeds, and they finally relax back to an isotropic state by the end of the process (Etournay et al., 2015; Iyer et al., 2019). Cell rearrangements are oriented along the AP direction initially before turning around to orient along the PD axis. We demonstrated that mechanical stress-dependent E-cadherin turnover influences cell shape changes and rearrangements during morphogenesis, which in turn determine the viscoelastic properties of the wing epithelium (Iyer et al., 2019). Most recently, we have shown that the core planar cell polarity pathway influences short-timescale cell mechanical properties without affecting the pattern of cellular dynamics or tissue mechanical stress (Piscitello-Gómez et al., 2022).

During the course of our work, we noticed that between the years 2016 and 2018, the onset of the process became consistently delayed. Here, we investigate the cause of this delay and we show that none of the tested factors can revert the delay in pupal wing morphogenesis. All fly strains tested during this work present the same delay, suggesting that there is a common factor that affects the delay in morphogenesis. Although *Drosophila* development has been shown to be sensitive to many environmental external factors such as temperature (Al-Saffar et al., 1996; Powsner, 1935) and diet (Alpatov, 1930; Ormerod et al., 2017; Reis, 2016), we find that the delay in pupal wing morphogenesis is unaffected by these factors. Although the cause of the delay remains unknown, our data hint at the relevance of light exposure. Our work provides novel insight into the robustness of pupal wing morphogenesis to environmental variation.

## 2 Results

### 2.1 Observation of a delay in the onset of pupal wing morphogenesis

When comparing our experiments on pupal wing morphogenesis conducted from 2011-2018, we have noticed that the onset of the process started to be consistently delayed sometime between 2016 and 2018. This delay is visible in the dynamics of cell elongation, which in 2011 reached a peak at approximately 22 h after puparium formation (hAPF) (Etournay et al., 2015) but in 2018 only by 28 hAPF (Iyer et al., 2019; Piscitello-Gómez et al., 2022). We highlight this observation in Fig 1. At 20 hAPF, cell shapes in the blade region are elongated in wings studied in 2011, 2012, and 2016, and they relax back to an isotropic shape by 28 hAPF (Fig 1A-B). In contrast, blade cells in flies studied after 2018 remain fairly isotropic until 22 hAPF but are highly anisotropic at 28 hAPF (Fig 1A-B). When we plot the entire dynamics of cell shape changes from timelapse imaging over the duration of pupal wing morphogenesis, we observe that the onset of morphogenesis in the 2018 flies is clearly delayed and the peak of cell elongation is achieved only by approximately 27-28 hAPF (Fig 1C).

**Figure 1:**
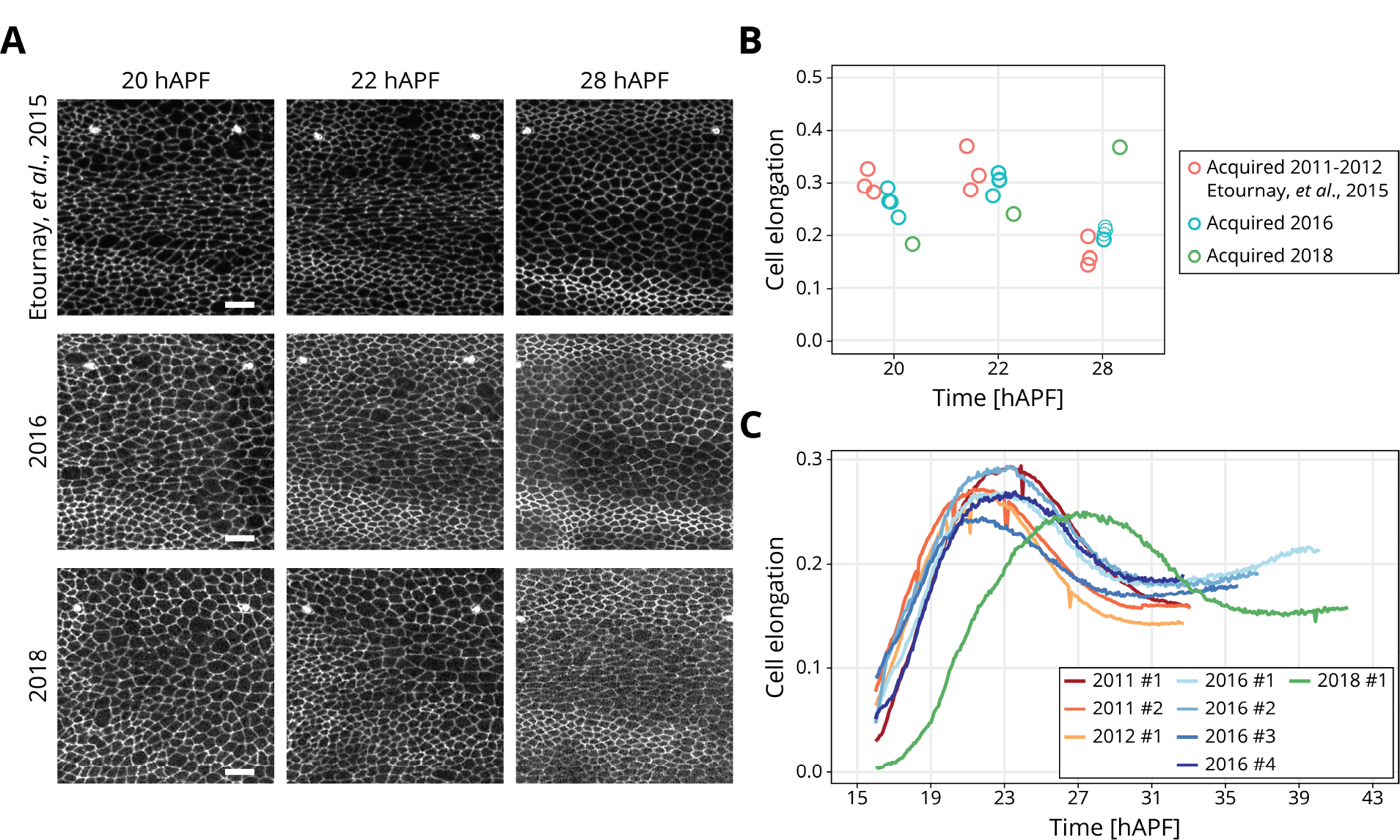
The peak of cell elongation is delayed in 2018 flies: (A) Snapshots of the blade region located between the second and third sensory organs in the intervein region between the L3 and L4 longitudinal veins at 20, 22 and 28 hAPF from three different long-term timelapses acquired in 2011-2012 (top row, published in Etournay et al., 2015), 2016 (middle row, Piscitello-Gómez et al., 2022) and 2018 (bottom row, Piscitello-Gómez et al., 2022). Scale bar, 10 *μ*m. (B) Quantification of cell elongation in a small region of the blade at 20, 22 and 28 hAPF from long-term timelapses acquired in 2011-2012 (red, n=3), 2016 (blue, n=4) and 2018 (green, n=1). (C) Complete dynamics of cell elongation during pupal wing morphogenesis for the whole blade of long-term timelapses acquired in 2011-2012 (red color palette, n=3), 2016 (blue color palette, n=4) and 2018 (green, n=1). Supplementary data: S1.1

We then plotted the rate of overall tissue shear and its decomposition into cell shape changes and cell rearrangements. We found that the rate of overall tissue deformation (blue curve in Fig S1.1A) is already high at the beginning of imaging in older flies, but in the 2018 flies, the rate of tissue shear is very low for the first 3 h. In addition, the timepoint when the direction of cell rearrangements (pink curve in Fig S1.1A) changes from from AP (negative shear rate) to PD oriented (positive shear rate) occurs 4-5 h later in 2018 flies than in 2011-2012 and 2016 flies (Fig S1.1). Similarly, the timepoint when the direction of cell elongation changes (from being PD-oriented, positive shear rate, to AP-oriented, negative shear rate), occurs 5 h later in 2018 flies than in 2011-2012 and 2016 flies (Fig S1.1). Lastly, the whole process ends by 32 hAPF in older flies, while it is still not completely finished until about 38 hAPF in the newer ones (Fig S1.1).

Note that from the outside, we can detect no visible difference in the age of the pupae. In our work, 16 hAPF has always been the starting point for imaging, because the pupa is very difficult to dissect any earlier. That continues to be the case now, so the age of pupa does not seem younger. Nonetheless, the delay of pupal morphogenesis relative to puparium formation now allows us to capture the process in its entirety, starting from when the rates of cell elongation changes, cell rearrangements, and tissue shear are close to zero. In contrast, in older experiments, the rates of tissue shear, cell elongation changes, and cell rearrangements were already high at the beginning of the experiment. Thus, it is difficult to directly compare the duration of the process to estimate the overall speed across the years. Nonetheless, if we instead estimate speed from part of process, for example from the time when the curves for cell elongation changes and cell rearrangements cross until the shear rates all return to zero, we see that the overall speed of the process is relatively unchanged (lines in Fig S1.1). Thus, we conclude instead that the major change across the years has to be a delay in the onset of the process, rather than a change in the overall speed of the process.

### 2.2 Imaging conditions do not account for the delay in onset of pupal wing morphogenesis

Our first thought was that the delay in pupal morphogenesis onset could be caused by imaging, as long-term timelapses of pupal wing morphogenesis involves a frequent exposure of the sample to light, which could also introduce subtle temperature fluctuations and phototoxicity. This explanation seemed unlikely, however, given that our settings for acquisition speed and power were always kept as constant as possible. Furthermore, our data from Fig S1.1 show clear differences in the rates of cell shape changes, cell rearrangements, and total shear at the beginning of imaging: with the post-2016 movies showing rates close to zero, whereas the pre-2016 movies showing high rates. These data suggest that the differences were present in the flies prior to the onset of imaging. Nonetheless, we sought to test directly how changing imaging parameters affects the timing of pupal wing morphogenesis. To simplify our analyses, we focused on measuring cell elongation magnitude in a small region of the blade: the L3-L4 intervein region between the second and third sensory organs.

We first investigated the effect of the repeated exposure to light. To do this, we analyzed cell elongation in *w*^-^; *Ecad::GFP;;* in movies where images were acquired either every 5 min or every 2 h at 25°C. While the peak of cell elongation in the movie with a time resolution of 5 min is reached at 28 hAPF (red curve in Fig 2), it occurs 2 h earlier in the movie with a time resolution of 2 h (green curve in Fig 2). Thus, more frequent imaging does slightly delay pupal morphogenesis. Nevertheless, the peak of cell elongation was not brought back all the way to 20-22 hAPF, where it was pre-2018 (Fig 2).

**Figure 2:**
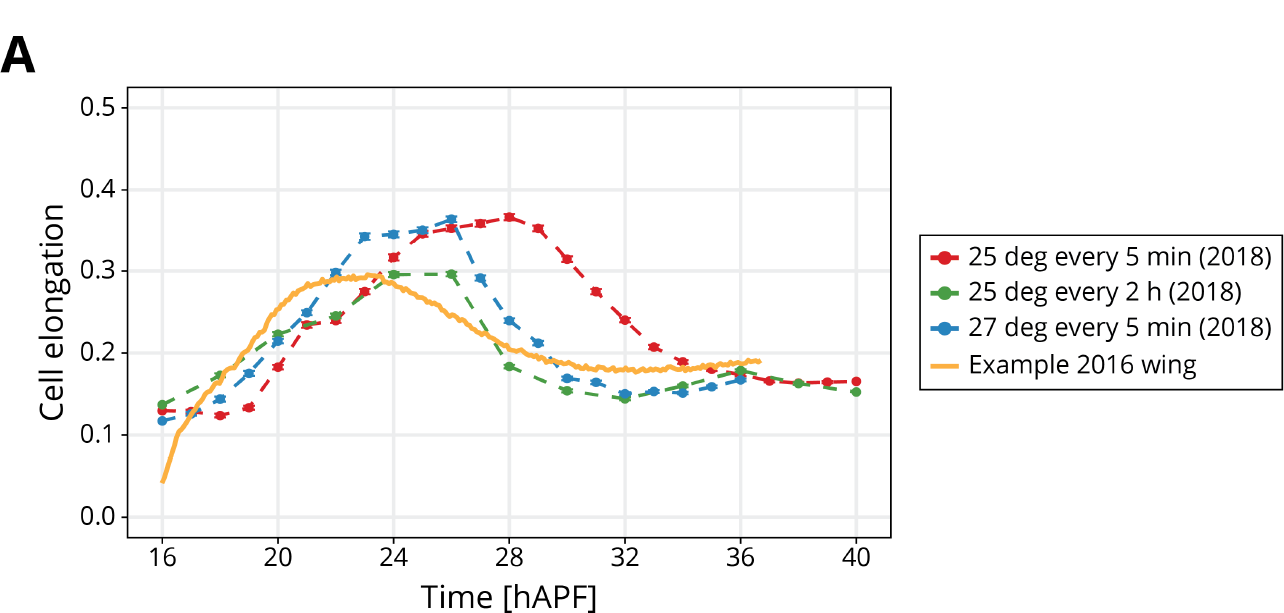
Imaging conditions have a minor effect on the timing of pupal wing morphogenesis: (A) Quantification of cell elongation magnitude in a small region of the blade throughout pupal wing morphogenesis for wings imaged every at 25°C every 5 min (red, n=1) or every 2 h (green, n=1), and at 27°C every 5 min (blue, n=1). The orange curve shows the cell elongation over time in one wing imaged in 2016. Filled circles show the average cell elongation for the region, and the line highlights the trend over time. The error bars correspond to the SEM.

We also tested the effect of temperature during image acquisition, reasoning that perhaps temperature control on the microscope had varied over the years or that the imaging itself may have caused subtle heating that would affect the process. To test for this possibility, we grew flies at 25°C but then switched the temperature on the microscope to 27°C and imaged every 5 min. We observe that the peak of cell elongation is reached at 26 hAPF (blue curve in Fig 2). Thus, increasing temperature can accelerate pupal morphogenesis slightly.

In conclusion, we observe that changes in acquisition rate and temperature can affect the timing of pupal wing morphogenesis, with increased light exposure slowing down and increased temperature speeding up the process. Nonetheless, even these relatively extreme changes in acquistion rate and temperature only shift the cell elongation curves by 2 h at most and therefore cannot account for the magnitude of shift in the onset of pupal wing morphogenesis that we have observed (5-6 h).

### 2.3 Changes in genetic background cannot account for the delay in pupal morphogenesis

Next, we wondered whether the delay in the onset of pupal wing morphogenesis is a consequence of a genetic change that had occurred in the line we were studying (*w*^-^; *Ecad::GFP;;*). To test for this possibility, we analyzed similar but distinct fly lines (Fig 3). Rather than acquiring full timelapse movies, we examined cell elongation in static timepoints, in order to greatly simplify the analysis and completely eliminate the effect of imaging conditions on the process. Specifically, we compared cell elongation at 20-22 hAPF (previous peak) to that at 28 hAPF (current peak). We also focused only on the small region of the blade used above. With this assay, we could rapidly determine whether the peak of cell elongation appeared to be shifted in the other lines as it was in *w*^-^; *Ecad::GFP;;*. The other lines tested include *w*^-^; *Ecad::GFP;;* and *w*^-^; *Ecad::GFP, pk*^*30*^;;, which contains a mutant in the planar cell polarity protein, Prickle, that does not seem to affect cell dynamics during this process (Piscitello-Gómez et al., 2022). Additionally, we outcrossed the *w*^-^; *Ecad::GFP;;* line for 4 generations to remove any potentially accumulated mutations (see Materials and Methods 5.5). All of these lines behave similarly to our original line, with higher cell elongation at 28 hAPF than at 20-22 hAPF, making it likely that the onset of pupal morphogenesis has shifted in all of these lines. We conclude that the genetic background is not relevant and that there may be an external environmental factor delaying the onset of pupal wing morphogenesis.

**Figure 3:**
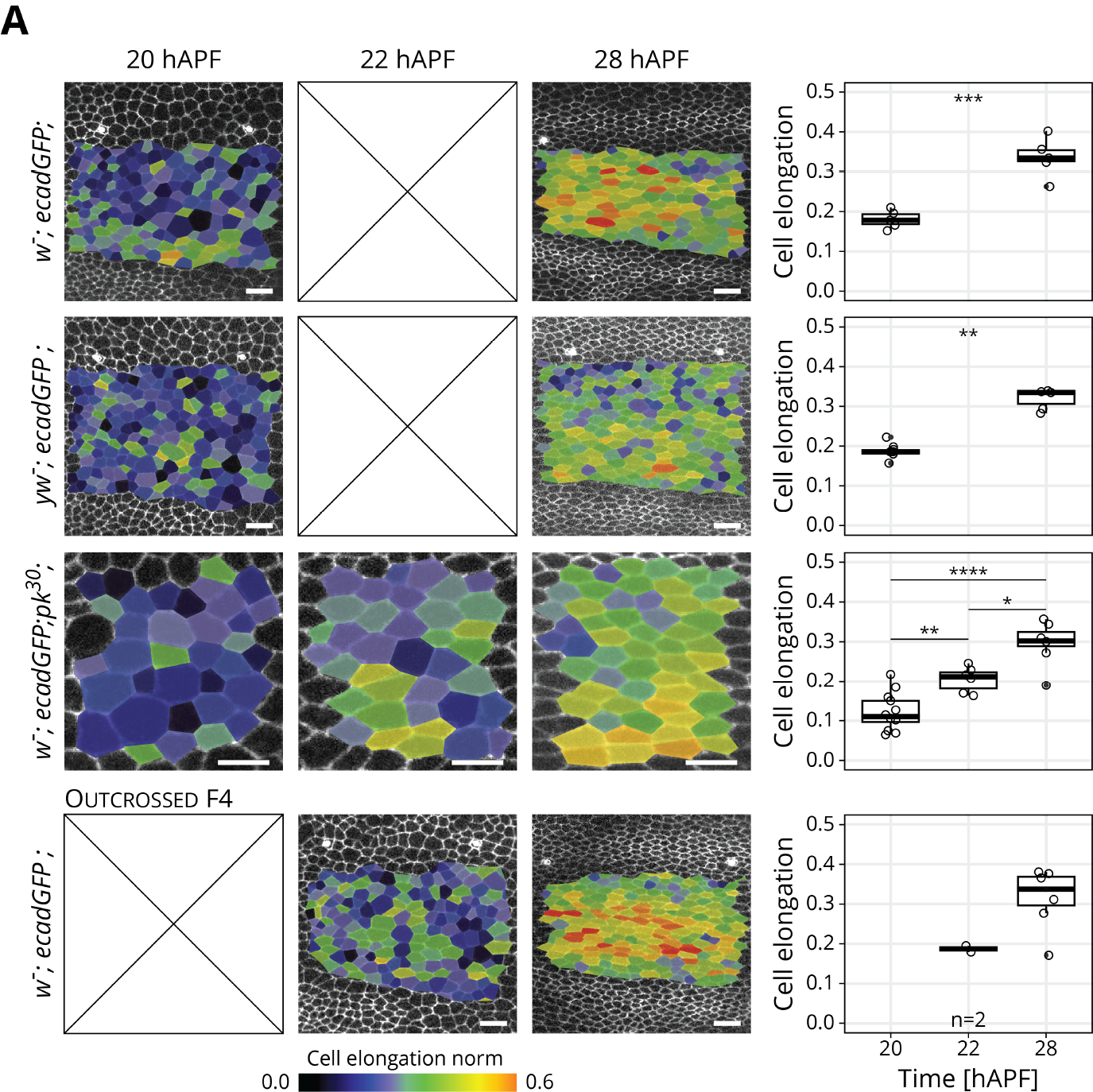
Changes in genetic background cannot account for the delay in pupal morphogenesis: (A) The images are shown for 20 or 22 hAPF and 28 hAPF. The fly lines studied were *w*^-*-*^; *Ecad::GFP;;*, first row), *w*^-^; *Ecad::GFP;;* (second row), *w*^-^; *Ecad::GFP, pk*^*30*^;; (third row) and *w*^-^; *Ecad::GFP;;* outcrossed for 4 generations (fourth row). Scale bar, 10 *μ*m, except for *pk*^*30*^: scale bar, 5 *μ*m. The corresponding quantification of the cell elongation magnitude in this region is show on the right column (n⩾3, except for the outcrossed line at 22 hAPF, where n=2). Note that the images and quantifications in *w*^*-*^; *Ecad::GFP, pk*^*30*^;; wings correspond to those of a smaller subregion of the blade region. Significance is estimated using the Kruskal–Wallis test. ***, p-val⩽0.001; **, p-val⩽0.01; ns, p-val*>*0.05. In all plots, each empty circle indicates one experiment, and the box plots summarize the data: thick black line indicates the median; the boxes enclose the 1st and 3rd quartiles; lines extend to the minimum and maximum without outliers, and filled circles mark outliers.

### 2.4 Timing of pupal morphogenesis is robust to many variations in rearing conditions

If not the imaging conditions, what other external or environmental factor(s) could cause a delay in pupal wing morphogenesis? To investigate this question, we sought to test on the impact on pupal morphogenesis of three factors that have been shown to affect overall *Drosophila* development: diet, parasitic infections, and light. As a readout, we again focused on cell elongation at 20 h vs 28 hAPF in *w*^-^; *Ecad::GFP;;* flies.

First, we investigated the effect of diet. Flies were fed with either yeast-or plant-based food, which differ in lipid composition and have been shown to affect *Drosophila* developmental rate and lifespan (Brankatschk et al., 2014, Brankatschk et al., 2018). We found that diet did not affect pupal morphogenesis, however: cell elongation was still significantly higher at 28 hAPF than at 20 hAPF (Fig 4A), with no significant differences between the two diets at either of these two different timepoints (Fig 4A). Thus, our data indicate that variations in the lipid content of larval diets does not alter the timing of pupal wing morphogenesis.

**Figure 4:**
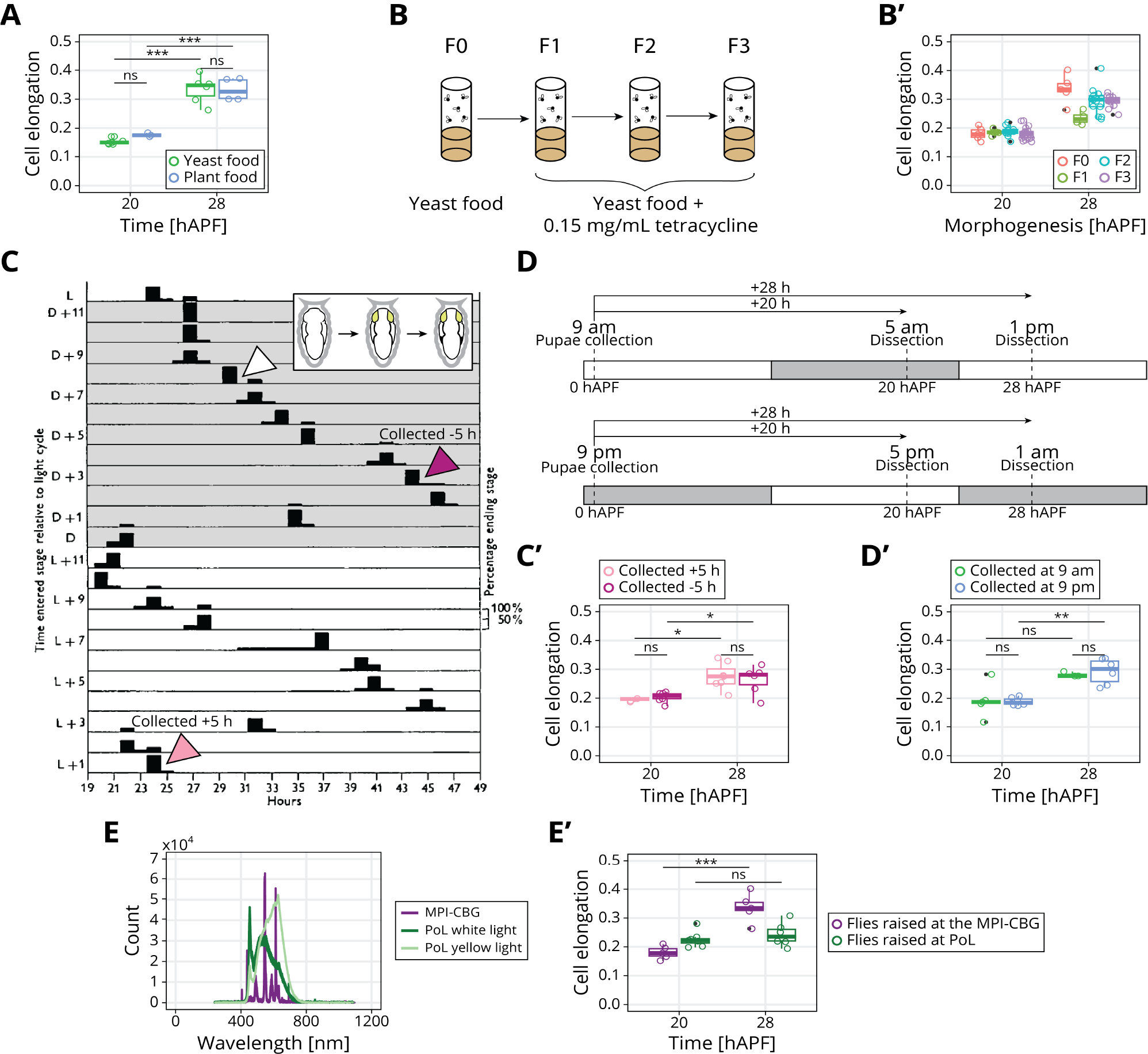
Diet, parasites, and light-darkness cycle do not affect pupal wing morphogenesis, but specific wavelengths of light exposure might: (A) Quantification of cell elongation magnitude in the blade region between the second and third sensory organs found in the intervein region between L3 and L4 in *w*^-^; *Ecad::GFP;;* flies that were raised on a yeast-based diet (green) or a plant-based diet (blue) (n*>*=3). (B) Tetracycline treatment in *w*^*-*^; *Ecad::GFP;;* flies over 3 generations. F0 was the initial untreated population, and the consecutive F1, F2, and F3 generations were treated with 0.15 mg/mL of tetracycline. (B^*’*^) Quantification of cell elongation magnitude in the blade subregion in *w*^-^; *Ecad::GFP;;* flies treated with tetracycline for 3 generations (n⩾4). (C) Graph from Harker, 1965. The plot shows the different durations of the time interval between head eversion and wing pigmentation (x-axis) depending on when in the light-darkness cycle head eversion occurs (y-axis). The gray area shows the hours when flies are in the dark, and the white area shows the hours when flies are exposed to light. The gray arrow indicates the duration of the time between head eversion and wing pigmentation in our usual collection, and the pink and purple arrows show the duration of this interval in pupae collected 5 h later (pink) or 5 h earlier (purple). (C^*’*^) Quantification of cell elongation norm in the blade subregion in pupae collected 5 h after (pink) and 5 h before (purple) the usual collection time (n⩾3). (D) Schematic of the experiment where pupae were collected 12 h apart and imaged at 20 and 28 hAPF. The imaging at 20 and 28 hAPF took place at opposing times in the light-darkness cycle. (D^*’*^) Quantification of cell elongation magnitude in the blade subregion in pupae collected at opposing times in the light-darkness cycle. Pupae collected while exposed to light (at 9 am) are shown in green, and those collected in the darkness are shown in blue (n⩾3). (E) Light spectra measured in the incubators where flies were raised in the MPI-CBG (purple) or PoL (dark green for measurements close to the white light, and bright green for measurements close to the yellow light). (E^*’*^) Quantification of cell elongation magnitude in the blade subregion in pupae raised in different light conditions (n⩾5). ***, p-val⩽0.001; **, p-val⩽0.01; *, p-val⩽0.05; ns, p-val*>*0.05. In all plots, each empty circle indicates one experiment, and the box plots summarize the data: thick black line indicates the median; the boxes enclose the 1st and 3rd quartiles; lines extend to the minimum and maximum without outliers, and filled circles mark outliers.

Second, we wondered whether the flies we had been working with had been infected with a parasite that could delay pupal wing morphogenesis onset. A common parasite that infects *Drosophila* and other arthropods is *Wolbachia* (Clark et al., 2005). Generally, the interactions between *Wolbachia* and *Drosophila* are symbiotic (Hamilton and Perlman, 2013), but in some cases *Wolbachia* has pathogenic effects on the host, such as feminization, male killing, and cytoplasmic incompatibility (Clark et al., 2005). To investigate whether this parasite was causing the observed delay, we first confirmed that flies were infected with *Wolbachia* by detecting its DNA in the flies (data not shown). Next, we eliminated the parasite from our *w*^-^; *ecadGFP;* flies by treating them with 0.15 mg/mL of tetracycline for 3 generations (Fig 4B). In the last generation (F3) *Wolbachia* DNA was no longer detected (data not shown), confirming that the treatment had worked. The cell elongation quantified in the blade subregion is still higher at 28 hAPF than at 20 hAPF in F3, however (Fig 4B_*′*_). Therefore, *Wolbachia* infection is not responsible for the observed delay.

Lastly, we studied the potential role of light in altering the timing of pupal wing morphogenesis. Specifically, we studied whether the timepoint of the light-dark cycle when pupae were collected played a role in the onset of pupal wing morphogenesis and therefore when the peak of cell elongation was reached. To do this, we performed two different kinds of experiments.

First, we performed similar experiments as those published in Harker, 1965. This study showed that the length of the time period between head eversion and wing pigmentation oscillates between 19 h and 47 h depending on when head eversion occurs in the light-dark cycle (Harker, 1965) (Fig 4C). For example, this interval can be as short as 24 h if head eversion occurs 1 h into the light phase (L+1, pink arrow in Fig 4C), while it extends to 44 h if head eversion occurs in the darkness (D+3, purple arrow in Fig 4C) (Harker, 1965). We therefore collected pupae at these extreme cases to test whether pupal wing morphogenesis onset could also be affected by the time when flies pupariated. Since the usual collection time for long-term time-lapses was 4 pm (head eversion occurs 12 h earlier at 4 am, which corresponds to D+8, white arrow in Fig 4D), we collected flies at 11 am (head eversion at 11 pm: D+3, dark magenta arrow in Fig 4D) and 9 pm (head eversion at 9 am: L+1, bright pink in Fig 4D). We did not observe significant differences between these two collections at neither 20 nor 28 hAPF. In both cases cell elongation was still higher at 28 hAPF than at 20 hAPF (Fig 4D*’*). From our data, we conclude that the timing of pupal wing morphogenesis does not depend on when in the light-dark cycle pupariation occurs.

Second, we collected pupae 12 h apart. One collection was done at 9 am, when the flies were exposed to light, and the other one was done in the darkness at 9 pm (Fig 4D-D_*′*_). With this approach wings were dissected at opposite times of the light-dark cycle (Fig 4D-D_*’*_). The quantification of cell elongation does not show significant differences between the flies collected at different day times, and cell elongation was higher at 28 hAPF than at 20 hAPF (Fig 4E_*’*_).

Lastly, we investigated whether the exposure to different light spectra influences the timing at which pupal wing morphogenesis occurs. In 2021, during the present study, our group moved from the Max Planck Institute of Cell Biology and Genetics (MPI-CBG) to the DFG Cluster of Excellence Physics of Life (shortened as PoL). These two institutions had different incubators for raising flies, which were each set to 25°C but used different light sources with differing wavelength spectra. The light in the MPI-CBG incubator was measured to have two big peaks around 550 and 600 nm (purple line in Fig 4E). In contrast, the incubator in PoL had two different light sources, classified as white and yellow light. Our spectra measurements close to the white light show that this light presents a big peak at 450 nm and a broader peak from 450 to 700 nm (dark green line in Fig 4E). The yellow light source, in contrast, is more shifted towards 600 nm (bright green line in Fig 4E). When raising flies in these two different light conditions, we observe that cell elongation is significantly higher at 28 hAPF than 20 hAPF in flies raised in the MPI-CBG. In PoL-raised flies, however, this difference between 20 h and 28 hAPF is no longer there, suggesting that the peak of cell elongation shifted. This result indicates that the spectra of light that the flies are exposed to during rearing can influence the timing of pupal wing morphogenesis.

### 3 Discussion/Summary

In this manuscript, we present the serendipitous observation that the start of pupal wing morphogenesis can be delayed by 5-6 h relative to the beginning of pupariation and we explore the potential genetic and environmental causes. This observation was made by comparing experiments performed over the course of nearly a decade in our lab: experiments performed before 2016 show morphogenesis starting significantly earlier than those performed after 2016. We show that it is really a delay in the onset of morphogenesis, as the complex choreography of cellular dynamics appears largely unchanged, proceeding at the same rate just shifted in time. We therefore then focus on cell elongation dynamics, comparing 20 hAPF (the previous peak of cell elongation) to 28 hAPF (the current peak of cell elongation). As this delay was observed in different genotype backgrounds, we conclude that it is caused by something in the environment. We then tested how many types of environmental variation influence the timing of pupal morphogenesis, including laser light illumination, temperature, diet, *Wolbachia* infection, and timing relative to the light-dark cycle. Somewhat surprisingly, we find that none of these can cause a 6 h shift in the peak of cell elongation. Therefore, this study reveals that the timing of pupal wing morphogenesis is actually fairly robust to environmental perturbation, making the original observation of a delay even more mysterious. Our data suggest that there may be an environmentally triggered gate controlling the onset of pupal wing morphogenesis; upon release, morphogenesis thereafter proceeds at a fairly constant rate. Although the nature of this environmental trigger remains unclear, it must be rather specific, as many sources of variation have no effect.

Importantly, in our screen for the environmental condition involved, we chose to greatly simplify our analysis, relying on the comparison of cell elongation in a particular region of the wing at 20 hAPF vs 28 hAPF, inspired by our original observation. There could, however, be more subtle/different effects of these environmental perturbations on pupal wing morphogenesis that were not captured with this method. By looking at the full timelapse of pupal wing morphogenesis, we indeed found that shifting temperature and illuminating light at different intervals does slightly affect the timing of morphogenesis, but the differences were relatively minor compared to our observation of a 6 h delay. It is an interesting observation in its own right that pupal morphogenesis proceeds largely unperturbed even in the face of these environmental alterations and the mechanisms maintaining such robustness are not clear.

Unfortunately, we were unable to identify the cause of the 6 h shift, although many factors can now be ruled out. One potential area to explore further is the light exposure. We found that a light sources emitting lower wavelengths produced flies with similar cell elongation magnitude at 28 hAPF as at 20 hAPF. A full timelapse analysis would reveal whether the peak of cell elongation is now shifted earlier or later, or whether there is now also an affect on overall rate. This observation, therefore, should motivate future experiments to clarify how and when different wavelengths of light may affect pupal wing morphogenesis or indeed any other events happening at the same time during pupal stages.

Lastly, our work has practical implications for future work on this process. Previously, we and others have timed pupal wing morphogenesis relative to the onset of pupariation. This work highlights a problem with this approach, as there seem to be cases in which pupal wing morphogenesis can be significantly delayed after the onset of pupariation. Thus, we recommend recalibrating time based on observations of the process itself. Specifically, we propose using the peak of cell elongation to recalibrate, reporting the peak as 0 h and then the rest of the timepoints relative to this point.

## 4 Supplementary Data

### 4.1 Fig 1 Supplementary

**Figure S1.**
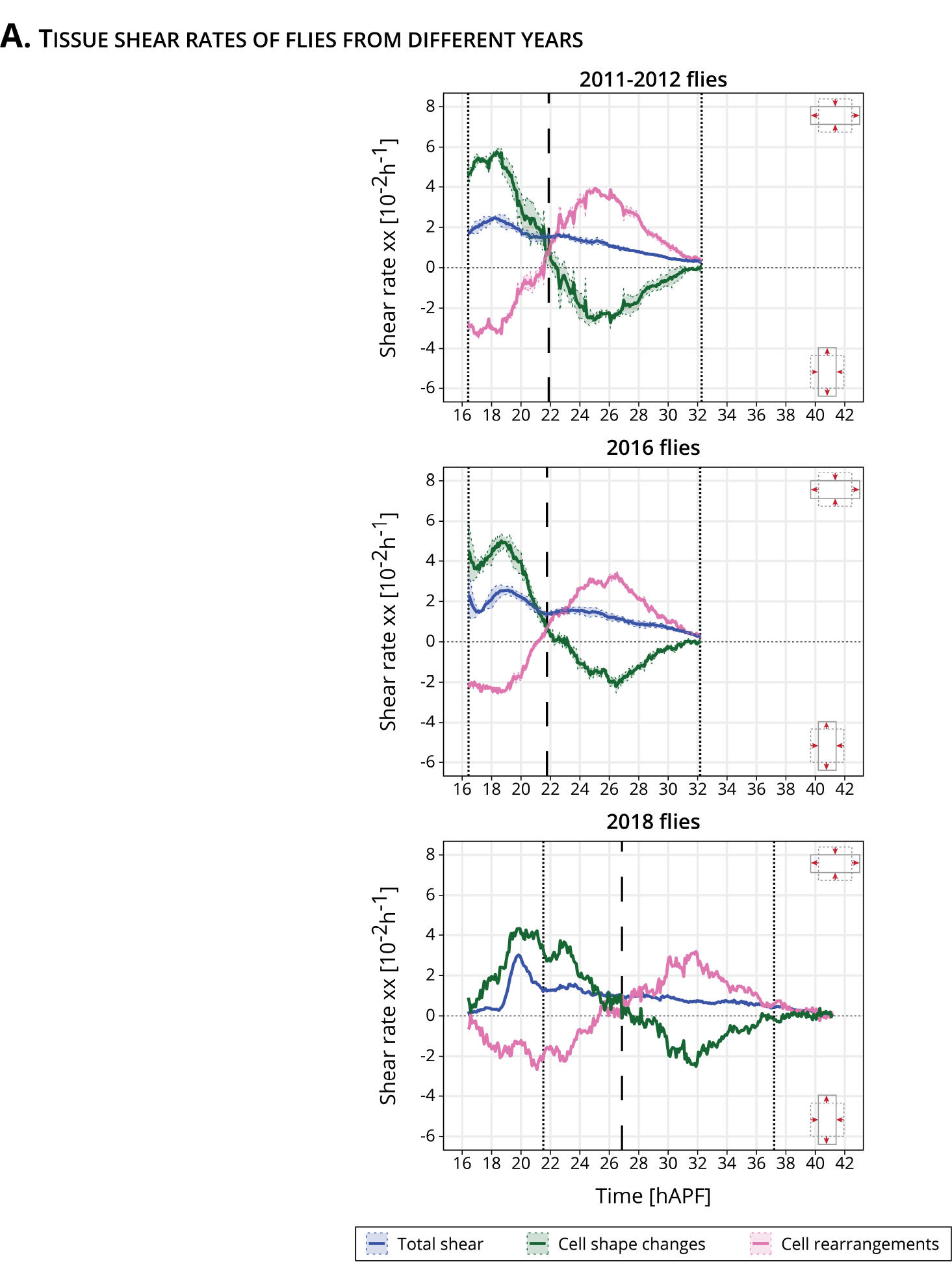
1: Tissue shear decomposition in *w*^-^; *Ecad::GFP;;* wings imaged in different years: (A) Cell dynamics underlying anisotropic blade deformation for 2011-2012 wings (n=3, top), 2016 flies (n=4, middle), and a 2018 fly (n=1, bottom). The dashed line highlights the timepoint when cell shape changes and cell rearrangements cross. The dotted lines highlight the beginning and end of the long-term timelapses in 2011-2012 and 2016 flies, and the interval between these two times is also shown in the last plot, corresponding to one timelapse of a 2018 fly.

## 5 Materials and Methods

### 5.1 Key resources table

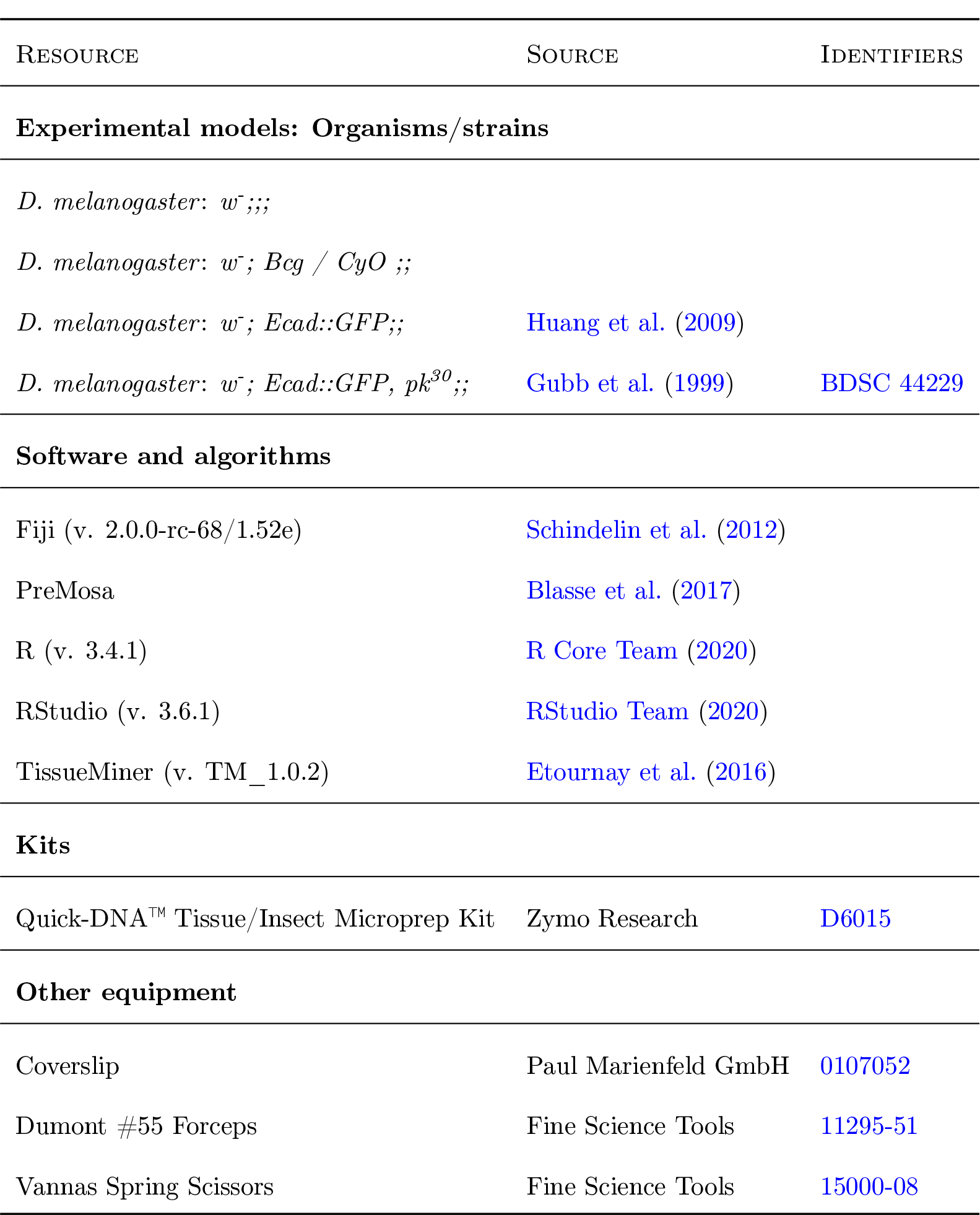

### 5.2 Fly husbandry

Flies were maintained at 25°C and fed with standard fly food containing cornmeal, yeast extract, soy flour, malt, agar, methyl 4-hydroxybenzoate, sugar beet syrup and propionic acid. Flies were kept at 25°C in a 12 h light/dark cycle, and they were flipped every 2-3 days. Experiments were performed with male flies, since their wings are smaller than female flies and require less tiles to be imaged.

### 5.3 Collection and imaging of pupal wings

White male pupa were collected and gently cleaned with a wet brush before being transferred to a vial with standard food. Flies were aged until the right developmental time (16 hAPF for long-term timelapses, and 20, 22 or 28 hAPF for snapshots). The pupal case was carefully dissected and mounted onto a 0.017 mm coverslip on a self-built metal dish with a drop of Holocarbon oil 700 (Classen et al., 2008).

#### 5.3.1 Long-term timelapse imaging of pupal wing morphogenesis

For long-term timelapses, pupae were collected at 4 pm, 5 pm and 6 pm in order to start the imaging at 8 am, 9 am or 10 am of the following day (when flies were 16 hAPF).

Long-term timelapses were acquired as described in Etournay et al., 2015 and Piscitello-Gómez et al., 2022. The microscope used was a Zeiss spinning disk microscope driven by ZEN 2.6 (blue edition), consisting of a motorized xyz stage, an inverted stand, a Yokogawa CSU-X1 scan head and a temperature-controlled chamber set to 25°C. The sample was illuminated with a 488 nm laser, and the emission was collected using a 470/40 bandpass filter. A Zeiss 63x 1.3 W/Glyc LCI Plan-Neofluar objective and a Zeiss AxioCam Monochrome CCD camera were used. The whole wing was imaged in 24 tiles with an 8% overlap. Each tile consisted of a Z-stack containing 50-60 slices separated by 1 *μ*m. Before imaging, the laser power was measured using a power-meter, and the exposure time and laser intensity were corrected to fulfill a laser power of 0.1 mW. An image was acquired either every 5 min or 2 h (for the 25°C every 2 h timelapse).

The long-term timelapse acquired at 27°C was acquired with an Olympus IX 83 inverted stand driven by the Andor iQ 3.6 software. The microscope is equipped with a motorized xyz stage, a Yokogawa CSU-W1 scan head and an Olympus 60x 1.3 Sil Plan SApo objective. The setup was located inside a temperature-controlled chamber set to 27°C. The sample was illuminated with a 488 nm laser, and the emission was collected using a 525/50 bandpass filter. The whole wing was imaged in 8 tiles with a 10% overlap. Each tile consisted of a Z-stack with 50-60 slices separated by 1 *μ*m. The laser power was set to 0.75 mW.

#### 5.3.2 Single snapshots

Single snapshots were acquired with the same Zeiss spinning disk microscope as described above but increasing the laser intensity and exposure time to 120 ms and 10.8%, respectively. In this case, only a small region of the blade was imaged in 9 tiles with a 8% overlap.

### 5.4 Processing, segmentation, tracking and database generation

Long-term timelapses, as well as single timepoints, were processed and analyzed as described in Piscitello-Gómez et al., 2022. Raw images were projected and stitched using using PreMosa (Blasse et al., 2017).

Long-term timelapses were registered using the Fiji plugin “Descriptor-based series registration (2D/3D + t)” and converted to 8 bit (Schindelin et al., 2012).

For both long-term timelapses and single snapshots, images were segmented using TissueAnalyzer (Aigouy et al., 2010; Aigouy et al., 2016). Segmentation errors were corrected manually by looking at the cell divisions and deaths masks. Images were rotated so that the angle formed by the line connecting the sensory organs would be 0. Regions of interest were manually defined. Lastly, a relational database was generated using TissueMiner (Etournay et al., 2016). We queried the information contained by the relational database using the Dockerized version of RStudio (Nickoloff, 2016).

#### 5.4.1 Cell elongation quantification

Prior to all laser ablation experiments, we acquired a 50 *μ*m stack that was projected using Premosa (Blasse et al., 2017). We cropped a region that enclosed the region that was ablated, segmented it using TissueAnalyzer (Aigouy et al., 2010; Aigouy et al., 2016) and generated a relational database with TissueMiner (Etournay et al., 2016).

The cell elongation measurements correspond to the norm of the cell elongation nematic tensor in the analyzed region. The definition of cell elongation was first presented in (Aigouy et al., 2010) and it describes the angle and magnitude of the tensor. The cell elongation tensor is given by

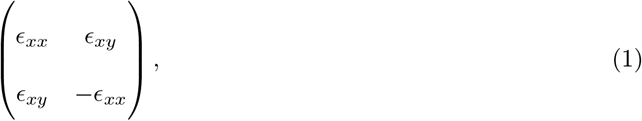

where

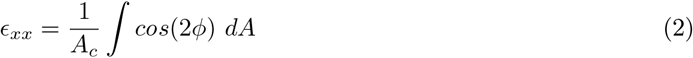

and

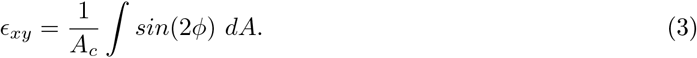

Cell elongation is normalized by the cell area (*A*_*c*_) of each cell. The magnitude of cell elongation is:

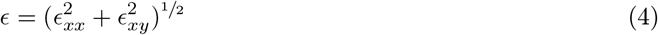

### 5.5 Outcross for 4 generations

The genetic background of the original *w*^-^; *Ecad::GFP;;* was cleaned up by outcrossing males with *w*^-^;;; virgins for 2 generations. The third generation was crossed with virgins *w*^-^; *Bcg / CyO ;;* to restore the homozygous *w*^-^; *Ecad::GFP;;* stock.

### 5.6 Experiments with different foods

Experiments with yeast and plant food were performed with food kindly provided by Dr. Suhrid Ghosh. The food composition was as follows:

- Yeast food: 80 g/L brewer’s yeast, 60 g/L glucose, 30 g/L sucrose, 20 g/L yeast extract, 20 g/L peptone and 10 g/L agar and 6 mL propionic acid (28.1% protein, 63.6% sugars and 8.2% lipids).
- Plant food: 160 g/L cornmeal, 76 g/L soy, 44 g/L treacle, 7 g/L agar, 2.61 g/L nipagen, 0.4 g/L vegetable oil and 12.6 mL propionic acid (26% protein, 68% sugars and 6% lipids).

Since flies do not lay many eggs in plant food, they were left in a cage on apple juice plates with a bit of yeast for 12 h to lay eggs. Afterwards, the plate was sprayed with PBS and the 100-150 *μ*L of eggs were transferred to vials with plant or yeast food and kept at 25°C.

### 5.7 Tetracycline treatment

*w*^-^; *Ecad::GFP;;* flies were tested for *Wolbachia* by isolating DNA from 10-20 male files using the Quick-DNA™ Tissue/Insect Microprep Kit. After DNA isolation, a PCR was run with the following *Wolbachia* specific primers (Jeyaprakash and Hoy, 2000):

The PCR product was run in a 1% agarose gel.

50 females and 50 males of the *w*^-^; *Ecad::GFP;;* stock were raised in vials with a concentration of 0.15 mg/mL of tetracycline for three generations. We confirmed the absence of *Wolbachia* in the third generation (F3) by PCR using the same protocol as above (data not shown). Positive and negative *D. melanogaster* controls were kindly provided by Wolfgang J. Miller, PhD (Lab Genome Dynamics, Center of Anatomy and Cell Biology, Medical University of Vienna).

**Table 1:**
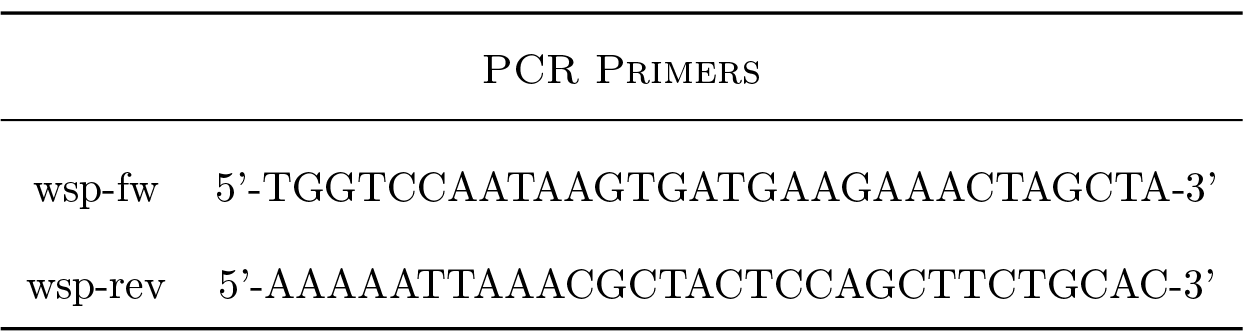
Wolbachia-specific PCR primers.

**Table 2:** PCR protocol.

| PROTOCOL |  |
| --- | --- |
| DNA | 1 μL |
| wsp-fw | 1 μL |
| wsp-rev | 1 μL |
| dNTPs | 1 μL |
| MgCl<sub>2</sub> | 0.6 μL |
| DMSO | 0.6 μL |
| Buffer | 4 μL |
| Polymerase | 0.5 μL |
| H<sub>2</sub>O | 10.3 μL |

### 5.8 Statistical analysis

Statistical analysis was done using RStudio (RStudio Team, 2020). We first always tested normality of the data using the Shapiro–Wilk test. If the data were normal, we used a Student’s t-test or ANOVA to test statistical significance between two or multiple groups, respectively. If the data were not normally distributed, significance was tested using the Mann-Whitney U test for two groups or Kruskal-Wallis test for multiple groups. Statistical test results are shown in the figure captions.

## Acknowledgements

We thank the Light Microscopy Facility, the Computer Department and the Fly Keepers of the MPI-CBG for their support and expertise. We also would like the thank Dr. Bert Nitzsche for his support measuring light spectra. *Wolbachia* positive and negative flies, as well as the sequence of the primers used for detection of *Wolbachia* DNA, were kindly shared by Prof. Wolfgang J. Miller from the Medical University of Vienna. We would like to thank Frank Jülicher, Marko Popovic and Jana Fuhrmann for their feedback and discussions. This work was funded by Germany’s Excellence Strategy – EXC-2068 – 390729961– Cluster of Excellence Physics of Life of TU Dresden, as well as grants awarded to SE from the Deutsche Forschungsgemeinschaft (SPP1782, EA4/10-1, EA4/10-2) and core funding of the Max-Planck Society to SE. NAD additionally acknowledges funding from the Deutsche Krebshilfe (MSNZ P2 Dresden). RPG was funded through the Elbe PhD program. We dedicate this work to our coauthor Prof. Dr. Suzanne Eaton, who tragically passed away before the finalization of the project.

## Author Contributions

**RPG**: Investigation, Formal analysis, Software, Validation, Data curation, Writing - original draft, Writing - Review and editing, Visualization. **AM**: Investigation, Writing - Review and editing. **NAD**: Resources, Supervision, Writing - original draft, Writing - Review and editing, Funding acquisition, Project administration. **SE**: Conceptualization, Methodology, Resources, Supervision, Funding acquisition, Project administration.

## Competing Interests

The authors declare no competing interests.

## Notes

### Competing Interest Statement

The authors have declared no competing interest.

